# The limits of metabolic heredity in protocells

**DOI:** 10.1101/2022.01.28.477904

**Authors:** Raquel Nunes Palmeira, Marco Colnaghi, Stuart A Harrison, Andrew Pomiankowski, Nick Lane

**Author notes:** **Corresponding author:** Nick Lane.

## Abstract

The universal core of metabolism could have emerged from thermodynamically favoured prebiotic pathways at the origin of life. Starting with H_2_ and CO_2_, the synthesis of amino acids and mixed fatty acids, which self-assemble into protocells, is favoured under warm anoxic conditions. Here we address whether it is possible for protocells to evolve greater metabolic complexity, through positive feedbacks involving nucleotide catalysis. Using mathematical simulations to model metabolic heredity in protocells, based on branch points in proto-metabolic flux, we show that nucleotide catalysis can indeed promote protocell growth. This outcome only occurs when nucleotides directly catalyse CO_2_ fixation. Strong nucleotide catalysis of other pathways (e.g. fatty acids, amino acids) generally unbalances metabolism and slows down protocell growth, and when there is competition between catalytic functions cell growth collapses. Autocatalysis of nucleotide synthesis can promote growth but only if nucleotides also catalyse CO_2_ fixation; autocatalysis alone leads to the accumulation of nucleotides at the expense of CO_2_ fixation and protocell growth rate. Our findings offer a new framework for the emergence of greater metabolic complexity, in which nucleotides catalyse broad-spectrum processes such as CO_2_ fixation, hydrogenation and phosphorylation important to the emergence of genetic heredity at the origin of life.

## Introduction

The problem of whether genes or metabolism arose first has a long and contested history (1-7). The question has limited value because nucleotide synthesis and polymerization necessarily require some form of protometabolism. A more useful approach is to investigate the protometabolic context in which genetic heredity first arose. One hypothesis for the origins of genetic heredity considers the emergence of RNA in protocells (8-10), where ‘autotrophic’ growth is driven by the continuous conversion of gases such as H_2_ and CO_2_ into organic matter via protometabolic pathways that prefigure the universal biosynthetic pathways (9). These are assumed to exist in the absence of genes or enzymes, meaning that the chemistry of life is thermodynamically and kinetically favoured (11, 12). Experimental work now links geochemical CO_2_ fixation with the universal core of metabolism in bacteria and archaea (13-15). Strikingly, core metabolic pathways including the acetyl CoA pathway (15, 16), large parts of the Krebs cycle (13, 17-19), glycolysis (20) and gluconeogenesis (21), the pentose phosphate pathway (14, 20) and some amino acid biosynthetic pathways (17, 22) occur spontaneously without enzymes. The idea that metabolism emerged from a geochemical protometabolism therefore looks increasingly persuasive (23-26).

The advantage of this hypothesis is that it readily explains how various components that make up protocells could come together to form a living system: the chemistry is driven *in situ* by a geologically sustained disequilibrium, primarily between H_2_ and CO_2_ (23, 27-30). But the idea that long, non-coded protometabolic pathways could arise spontaneously and be reliable enough to foster the emergence of the genetic code seems to strain credibility. In particular, nucleotide synthesis is missing from the pathways listed above. Purine nucleotide synthesis requires twelve steps, albeit with some repetitive chemistry, beginning with amino acids, sugar phosphates and an energy currency. There have only been limited experimental attempts to synthesise nucleotides prebiotically following these biological pathways (9, 31). Yet even if they are successful, could protometabolism really generate enough nucleotides to form RNA?

If not, even the best-case scenario falls short of the requirements for the emergence of genetic heredity in autotrophic protocells. Using life as a guide to its own origin is then either misguided, or it must be possible for protocells to get better at making nucleotides before the emergence of genetic heredity. The simplest possibility is that nucleotides can catalyse their own synthesis, either directly or indirectly. Nucleotide-derived cofactors include coenzyme A, NADH, FADH, ATP, and the pterins and folates involved in CO_2_ fixation (32-34). ‘Naked’ cofactors frequently catalyse the same chemistry as the holoenzymes, albeit more slowly (35-37). It is therefore possible to imagine that nucleotides could form at trace levels within protocells and favour protocell growth and replication through simple catalysis. But could such catalysis increase nucleotide concentrations as well as protocell growth? If so, would the amplification of nucleotide synthesis in protocells depend on autocatalysis (nucleotide catalysis of nucleotide synthesis) or positive feedbacks at the network (protocell) level?

Theoretical models allow a quantitative analysis of what is possible, given the architecture of the system. We have previously shown that positive feedbacks involving FeS clusters chelated by amino acids can drive the growth of simple protocells composed of fatty acids and amino acids through CO_2_ fixation (38). Experimental work confirmed two key predictions: robust protocells do assemble under the conditions proposed (39, 40), and ‘biological’ FeS clusters form spontaneously in the presence of monomeric amino acids such as cysteine (41). Here, we develop the protocell model to consider the synthesis of nucleotides via a branching protometabolism based on universally conserved pathways (42-45). The model addresses how alterations in protometabolic flux affect protocell growth. Catalysis lowers the barriers to specific reactions, diverting flux down particular pathways (46). But increased flux down one pathway (e.g. towards fatty acids) diminishes flux down other pathways (e.g. towards amino acids). We consider the full network of interactions to evaluate how nucleotide catalysis could contribute to protocell growth. These include nucleotide catalysis of CO_2_ fixation and specific protometabolic pathways (23, 29, 47-49)) as well as nucleotide autocatalysis, and the cascade of feedbacks that ensue. The model reveals the necessary architecture of nucleotide catalysis that favours growth, and examines whether such processes could generate enough nucleotides to facilitate polymerization and the emergence of genetic heredity.

### Model overview

We develop an earlier model of protocell metabolism grounded in prebiotic chemistry as the basis for our approach here (38). The model assumes that the ions Fe^3+^ and S^2–^ are chelated by monomeric amino acids to form FeS clusters within protocells. These clusters associate with the membrane, where they draw on geochemically sustained proton gradients to facilitate CO_2_ fixation (29) (figure 1*a*). The first steps of CO_2_ fixation form a two-carbon prebiotic equivalent to acetyl CoA (e.g. methylthioacetate (23, 48, 50)) here labelled *C*_2_. We assume that Fe^3+^ and S^2–^ ions are not rate-limiting, so the rate of CO_2_ fixation is proportional to the number of amino acids capable of forming FeS clusters present in the cell.

**Figure 1.**
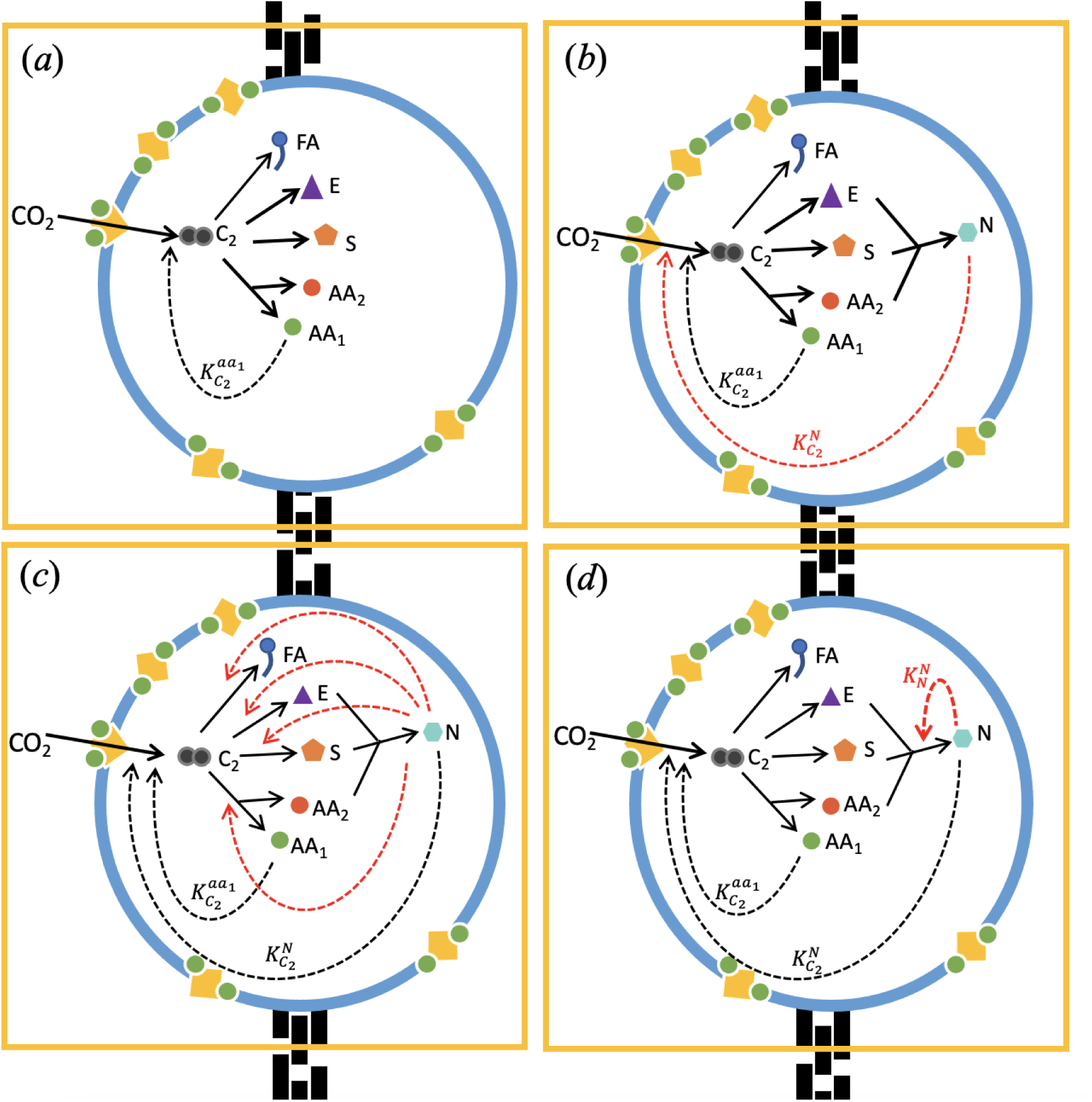
Models of positive feedbacks in protocells. Each panel depicts protocell models of autotrophic CO_2_ fixation driven by H_2_ (not shown) and catalysed by membrane-bound iron-sulphur clusters chelated by amino acids (yellow squares with green circles) under hydrothermal-type conditions. Fixed CO_2_ initially forms a simple two-carbon activated acetate, equivalent to acetyl CoA (C_2_), which acts as the primary substrate for a branching protometabolism based on the universally conserved core of biochemistry. (a) The null model considers the base-case in the absence of nucleotide formation in which the products are fatty acids (FA), amino acids, half of which (AA_1_) feedback on CO_2_ fixation, sugars (S) and energy (E). The subsequent models consider the catalytic function of nucleotides (N) produced from sugars (S), energy (E) and the other half of amino acids (AA_2_). In (b) nucleotides catalyse CO_2_ fixation 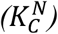 alone; in (c) nucleotides catalyse individual synthetic pathways (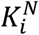, where i ∈ FA, AA_1_, AA_2_, S, E) in addition to CO_2_ fixation; and in (d) nucleotides perform autocatalysis 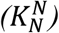 in addition to CO_2_ fixation. Solid arrows represent synthesis reactions and dotted arrows represent catalysis and are shown in red for the variant models. Note that the energy currency E (equivalent to acetyl phosphate) is consumed by nucleotide synthesis, hence is equivalent to a substrate and is shown by a solid line.

The model is developed by introducing several metabolic branch points that result in *C*_2_ being converted into amino acids (*AA*), fatty acids (*FA*), sugars (*S*) and a primitive energy currency (*E*) (e.g. acetyl-phosphate (48); figure 1*b*). We do not model individual steps but assume that whole pathways are capable of arising spontaneously (51). Other organic molecules would be produced from *C*_2_, but these are not considered further as they are irrelevant to turnover. The production of two types of amino acids is considered. The first set, *AA*_1_, chelates FeS clusters, which partition to the membrane and catalyse CO_2_ fixation. The second set, *AA*_2_, combines with sugars and energy currency to produce nucleotides (figure 1*b*). The proportion of *C*_2_ channelled into the synthesis of each species is inferred from thermodynamic data under alkaline hydrothermal conditions ((11), Table 1), which favours production of amino acids and fatty acids (56.5% and 37.5% respectively) over sugars (5%), whereas energy production is set at a minimal level (1%) corresponding to low availability of inorganic phosphate (52-56). As the amount of fixed carbon available is finite, increasing the production of one species decreases the production of others. The model also describes protocell division, which occurs when the cell surface area (proportional to the fatty-acid content) reaches a critical size. The daughter protocell inherits half the number of molecules of all species in the protocell.

**Table 1.** Main parameters used for simulations.

| Parameter | Symbol | Value | Unit |
| --- | --- | --- | --- |
| Rate constant for catalysis of CO<sub>2</sub> fixation by nucleotides | $K_C^N$ | 10<sup>3</sup> | mol<sup>-1</sup> dm<sup>3</sup> |
| Rate constant for catalysis of nucleotide production by E | $K_N^E$ | 2 | mol<sup>-3</sup> dm<sup>9</sup> s<sup>-1</sup> |
| Rate constant for catalysis of nucleotide production by nucleotide cofactors | $K_N^N$ | 2 | mol<sup>-1</sup> dm<sup>3</sup> |
| Saturating constant for catalysis of nucleotide production by nucleotides | $K_i^N$ | 1 | cm<sup>-1</sup> s<sup>-1</sup> |
| Proportion of C<sub>2</sub> molecules used to produce fatty acids | $\lambda_{FA}^{C_2}$ | 0.376 | Unitless |
| Proportion of C<sub>2</sub> molecules used to produce amino acids | $\lambda_{AA}^{C_2}$ | 0.564 (0.282 into each type of amino acid) | Unitless |
| Proportion of C<sub>2</sub> molecules used to produce the energy currency (E) | $\lambda_E^{C_2}$ | 0.01 | Unitless |
| Proportion of C<sub>2</sub> molecules used to produce sugars | $\lambda_S^{C_2}$ | 0.05 | Unitless |

The dynamics of protocell metabolism are modelled by a system of ordinary differential equations describing the change in the number of molecules of each species over time. These changes are determined by the number of stoichiometric reagents and catalysts present in the system. Besides amino-acid catalysis of CO_2_ fixation, which is a feature of the null model (figure 1*a*), three additional catalytic positive feedbacks are considered: (1) catalysis of CO_2_ fixation by nucleotides (figure 1*b*); (2) catalysis of individual protometabolic pathways alongside CO_2_ fixation (figure 1*c*); and (3) nucleotide autocatalysis alongside CO_2_ fixation (figure 1*d*). A full description of the model is given in the SI.

## Results

### Nucleotide catalysis of CO_2_ fixation

The model assumes that CO_2_ fixation produces fatty acids (*FA*), which directly contribute to membrane growth, and amino acids, some of which chelate FeS clusters (*AA*_1_), providing a positive feedback loop that enhances CO_2_ fixation (figure 1*a*). Additional branch points generate further species (*AA*_2_, *S* and *E*), which contribute to nucleotide production (figure 1*b*). In the absence of nucleotide production (figure 1*a*) or nucleotide catalysis (i.e.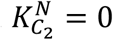, figure 1*b*), this has no effect on cell growth. However, the additional metabolic branch points pay back when nucleotides catalyse CO_2_ fixation (i.e 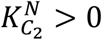; figure 1*b*). Nucleotide catalysis of CO_2_ fixation 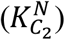 generates a positive feedback, increasing the rate of cell division as the strength of catalysis increases (figure 2).

**Figure 2.**
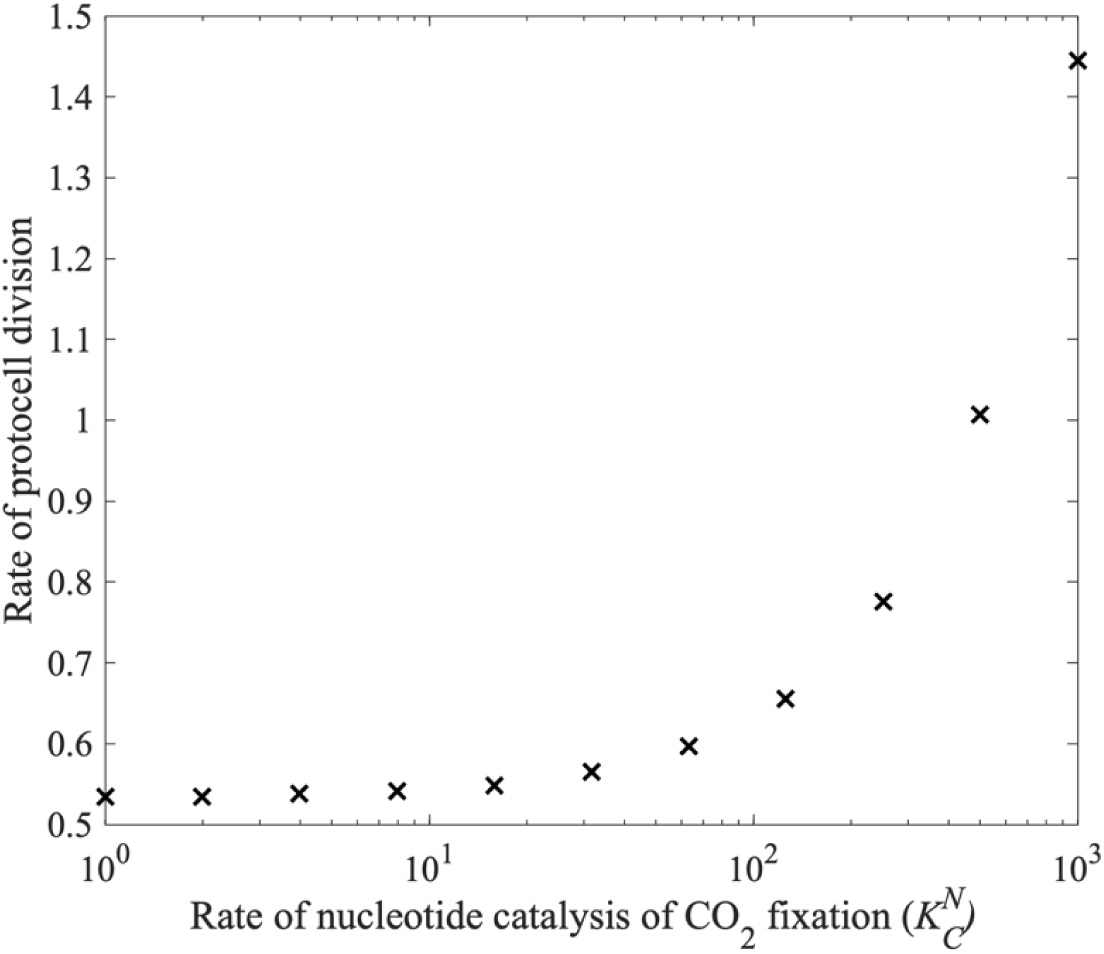
The effect of nucleotide catalysis of CO_2_ fixation 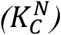 on the rate of cell division. Other parameter values given in Table 1.

### Nucleotide catalysis of individual pathways

Taking catalysis of CO_2_ fixation as the base-case for comparison (figure 1*b*), we consider the consequences of nucleotide catalysis of individual branch points, enhancing the production of particular species in protocells (figure 1*c*). These consequences are observed through changes to the rate of protocell division (figure 3) and nucleotide concentration within protocells (figure 4).

**Figure 3.**
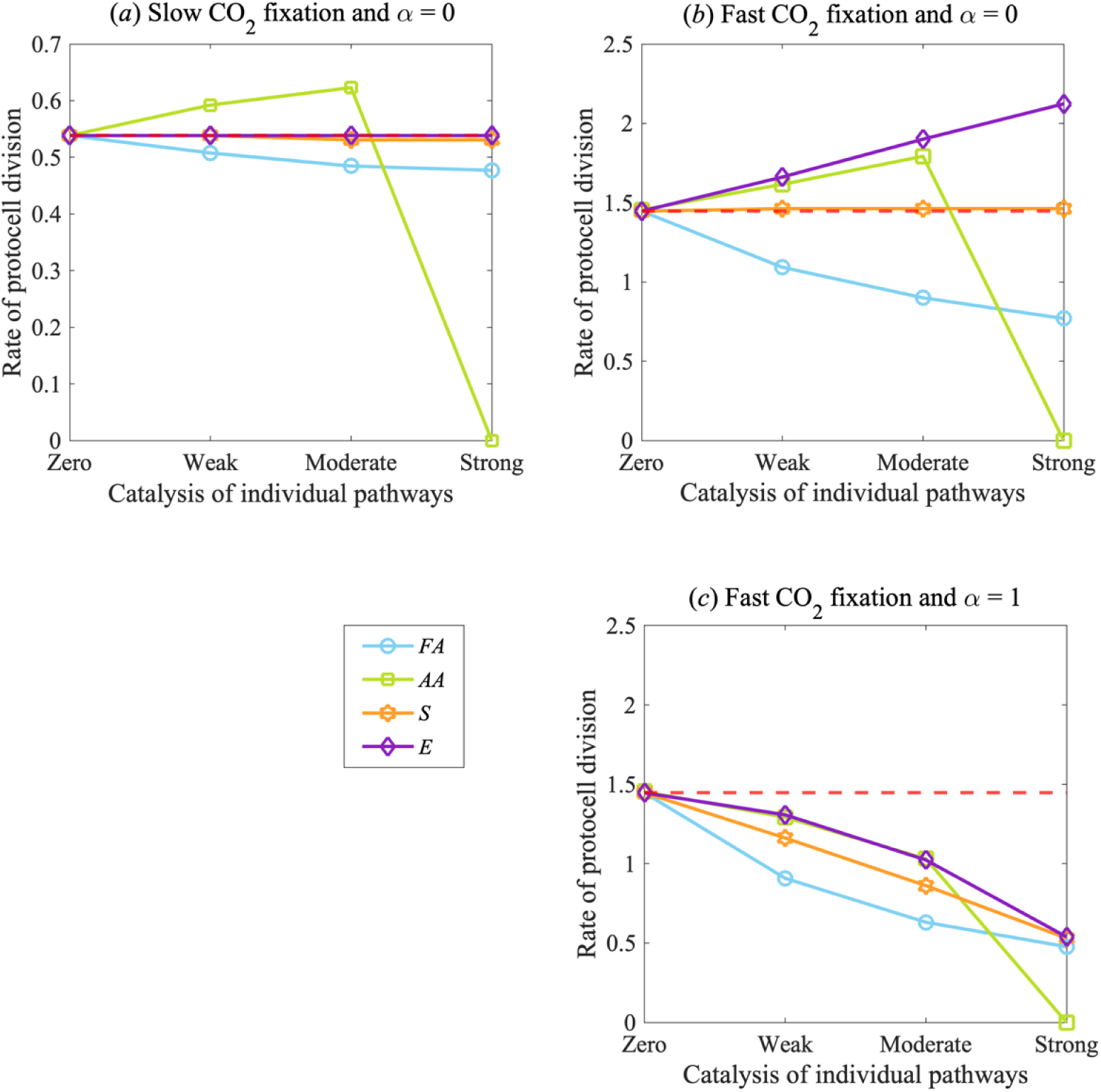
Impact of nucleotide catalysis of individual pathways on protocell division rate. Changes in protocell division per day is shown for nucleotide catalysis of the synthesis pathways of amino acids (AA, green), fatty acids (FA, blue), energy (E, purple) and sugars (S, orange). In each panel, nucleotide catalysis ranges from zero 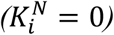 to weak 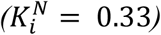, moderate 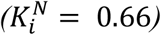 and strong 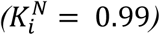 of specific pathways i = AA, FA, S, E. These rates are considered in relation to (a) slow 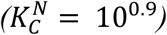 and (b) fast 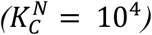 rates of CO_2_ fixation, given no cost to catalysis (α = 0). In (c) a cost to catalysis (α = 1) is introduced in proportion to the strength of nucleotide catalysis 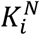 (with fast rates of CO_2_ fixation). The red dotted line indicates the number of protocell divisions per day with nucleotide catalysis of CO_2_ fixation alone (i.e. no catalysis of other synthesis pathways, as in figure 1b). Other parameter values given in Table 1.

**Figure 4.**
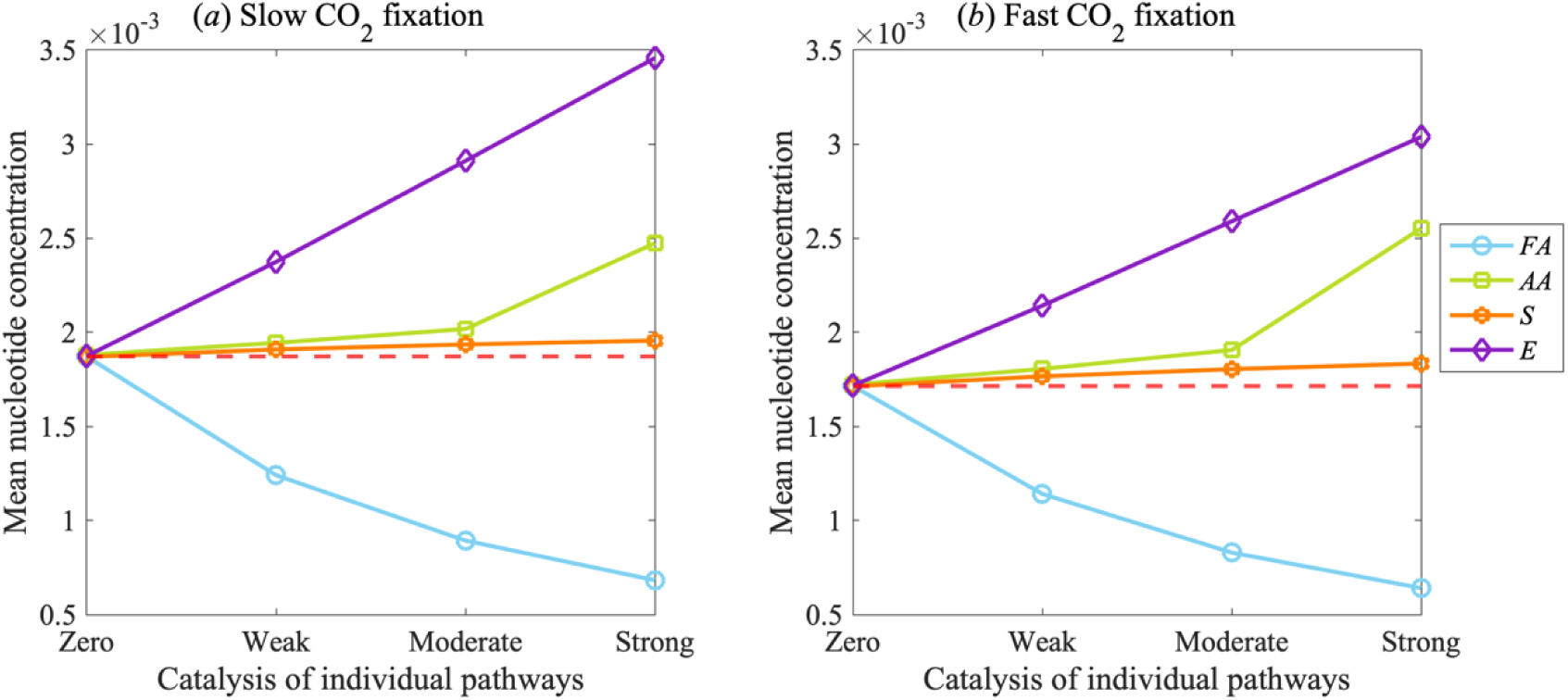
Nucleotide accumulation depending on catalysis of individual pathways. Changes in mean nucleotide concentration at equilibrium with increasing rates of nucleotide catalysis of specific pathways, using the same values as figure 3. (a) slow 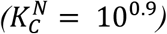 or (b) fast 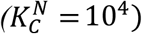 rates of CO_2_ fixation, assuming no cost to catalysis (α = 0). The red dotted line indicates the number of protocell divisions per day with nucleotide catalysis of CO_2_ fixation alone (i.e. no catalysis of other synthesis pathways). Other parameter values are given in Table 1.

Nucleotide catalysis of amino-acid synthesis 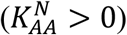 leads to a small increase in the rate of protocell division when the strength of 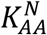 is weak or moderate (figure 3*a*). This is because increased amino-acid synthesis (*AA*_1_) forms more FeS clusters, which promote CO_2_ fixation and so protocell growth through the positive feedback loop on amino-acid synthesis. This positive-feedback loop also generates more of the second set of amino acids (*AA*_2_), which contribute to nucleotide production, potentially enhancing the catalytic feedback of nucleotides on CO_2_ fixation. But that effect is self-limiting, as it does not overcome the disadvantage of channelling carbon away from other species. This disadvantage is clearly seen when catalysis increases the rate of the production of only one type of amino acid, leaving the other type unchanged (figure S1). For these reasons, protocell growth and division collapses with strong catalysis of amino acids (figure 3*a*). If fixed carbon is excessively channelled into amino-acid synthesis (when 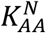 strength is strong), that precludes fatty-acid synthesis and inhibits growth and division. These results are independent of the strength of nucleotide catalysis on CO_2_ fixation. With fast nucleotide catalysis of CO_2_ fixation, the rate of protocell division increases, but the same collapse occurs with strong catalysis of amino-acid synthesis for the same reasons – flux is diverted towards amino acids and away from fatty acids, which are necessary for protocell growth (figure 3*b*).

In contrast, if nucleotides catalyse fatty-acid synthesis 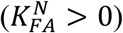, this invariably undermines protocell division (figure 3*a*-*b*). Even though fatty acids directly contribute to protocell growth (by increasing membrane surface area), there is no positive feedback loop associated with fatty-acid synthesis. As a result, the advantage to protocell growth of channelling *C*_2_ into the production of fatty acids does not overcome the disadvantage of channelling *C*_2_ away from other products. Again, these results are independent of the catalytic strength of nucleotides on CO_2_ fixation, but are more dramatic with higher 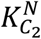 as this diverts a greater proportion of flux towards fatty acid synthesis and away from other useful products (figure 3b).

Catalysis of the pathways that synthesise an energy currency 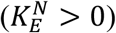 or sugars 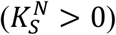 has a neutral or mildly deleterious effect on protocell growth when nucleotides catalyse CO_2_ fixation only weakly (figure 4*a*). The proportion of *C*_2_ used in the production of energy and sugars is too low to channel much fixed carbon away from the production of amino acids and fatty acids, even with strong catalysis of these pathways. But if nucleotide catalysis strongly favours CO_2_ fixation, there is a benefit to channelling more resources towards an energy currency, which becomes more apparent as 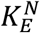 increases (figure 4*b*). This difference reflects the assumption that less fixed carbon is channelled towards an energy currency than to sugar synthesis (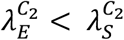, Table 1). Thus, making more energy currency helps because this species limits the production of nucleotides. Sugars are not limiting, so increasing their synthesis does not substantially influence nucleotide production. This interpretation is corroborated if we assume that more carbon is needed to make sugars (and so fewer sugar molecules are made per unit of *C*_2_) (figure S2). In this case, the availability of sugars begins to limit nucleotide synthesis, so strong catalysis of sugar production now increases protocell growth.

### Concentration of nucleotides

As well as altering the rate of protocell division (figure 3), nucleotide catalysis of individual pathways can alter the concentration of nucleotides within the protocell (figure 4). Nucleotide concentration only increases markedly when nucleotides catalyse the production of an energy currency 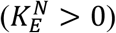. Again, this arises because energy is the limiting factor for nucleotide synthesis, so increasing the synthesis of an energy currency feeds through to a linear increase in nucleotide concentration within the protocell (figure 4*a*). That contrasts with catalysis of sugar production 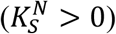, which has at best a slightly positive effect on nucleotide concentration (figure 4*a*). The difference reflects the thermodynamic assumption that sugars are formed in excess of the requirements for nucleotide synthesis, so increasing their production does not substantially influence the concentration of nucleotides in the protocell.

Nucleotide catalysis of amino-acid synthesis 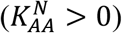 likewise has a marginal effect on nucleotide concentration, as channelling *C*_2_ towards amino acids retards the synthesis of sugars and an energy currency, whose production is essential to nucleotide generation (figure 4*a*). In addition, because weak or moderate amino-acid catalysis of CO_2_ fixation drives protocell growth and division (figure 3*a*), nucleotide concentration is halved more regularly with each cell division, lowering their steady-state concentration. In contrast, strong catalysis of amino-acid synthesis favours higher concentrations of nucleotides (figure 4*a*) because this collapses protocell growth (figure 3), allowing nucleotides to accumulate, while also synthesising the amino acids needed for nucleotide synthesis (*AA*_2_) (figure 3*a*). This latter point explains the difference with fatty-acid synthesis. Nucleotide catalysis of fatty-acid synthesis 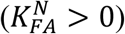 not only reduces protocell division but also consistently lowers the concentration of nucleotides in protocells (figure 4*a*). That is because channelling *C*_2_ away from the precursors of nucleotide synthesis (amino acids, sugars and an energy currency) leads to a progressive decrease in concentration of nucleotides.

These qualitative results are largely independent of the catalytic effect of nucleotides on CO_2_ fixation (figure 4). Faster nucleotide catalysis of CO_2_ fixation still generates similar levels of nucleotides in protocells (figure 4b). That is because fast catalysis of CO_2_ fixation 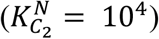 not only increases the rate of nucleotide production but also the rate of protocell growth and division, leaving nucleotide concentration at roughly the same level. A minor exception to this pattern is seen when nucleotides catalyse energy production, which leads to a slight decrease in the concentration of nucleotides with faster catalysis of CO_2_ fixation (i.e. figure 4*b* vs figure 4*a*, purple lines). In this case, the energy currency drives nucleotides synthesis, which produces a stronger feedback on cell growth and division that in turn depletes nucleotide concentrations.

### Costs of additional catalytic pathways

For the results described above, there is no cost to CO_2_ fixation through the introduction of additional catalysis of individual pathways (*α* = 0). This lack of cost is consistent with one type of nucleotide catalysing CO_2_ fixation and different types catalysing specific pathways. An alternative modelling possibility is that increased catalysis of specific pathways decreases the catalysis of CO_2_ fixation. This would be the case if the same type of nucleotide performed both catalytic functions (*α* > 0). In this case, catalysis of individual pathways leads to a decrease in the rate of CO_2_ fixation, which feeds through to a decrease in the rate of protocell division (figure 3*c*). This effect applies whichever metabolic branch point is catalysed: the stronger the competitive catalysis of specific pathways, the greater the negative effect on CO_2_ fixation and the corollary decline in protocell growth rate (figure 3*c*).

### Nucleotide autocatalysis

A final possibility analysed is direct nucleotide catalysis of their own synthesis, i.e. direct rather than network autocatalysis (figure 1*d*). This can be advantageous for protocell growth when autocatalysis of nucleotides and catalysis of CO_2_ fixation are coupled and both 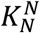 and 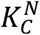 are high (figure 5*a*). However, this combination does not generate high concentrations of nucleotides in the protocell (figure 5*b*) for similar reasons to those noted above. Although nucleotides are produced at a faster rate, they do not accumulate because with a high rate of CO_2_ fixation their production is coupled to an increased rate of protocell growth and division. Nucleotide accumulation is greatest when autocatalysis 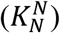 is high but catalysis of CO_2_ fixation 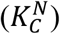 is low (figure 5*b*), which undermines the rate of protocell division (figure 5*a*) allowing nucleotide accumulation (figure 5*b*). A negative correlation between nucleotide autocatalysis 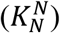 and catalysis of CO_2_ fixation 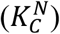 is observed if there is a cost interaction between the two types of catalytic pathways, meaning that the highest nucleotide concentrations are only generated at the cost of protocell division and vice versa (figure 5*a*-*b*).

**Figure 5.**
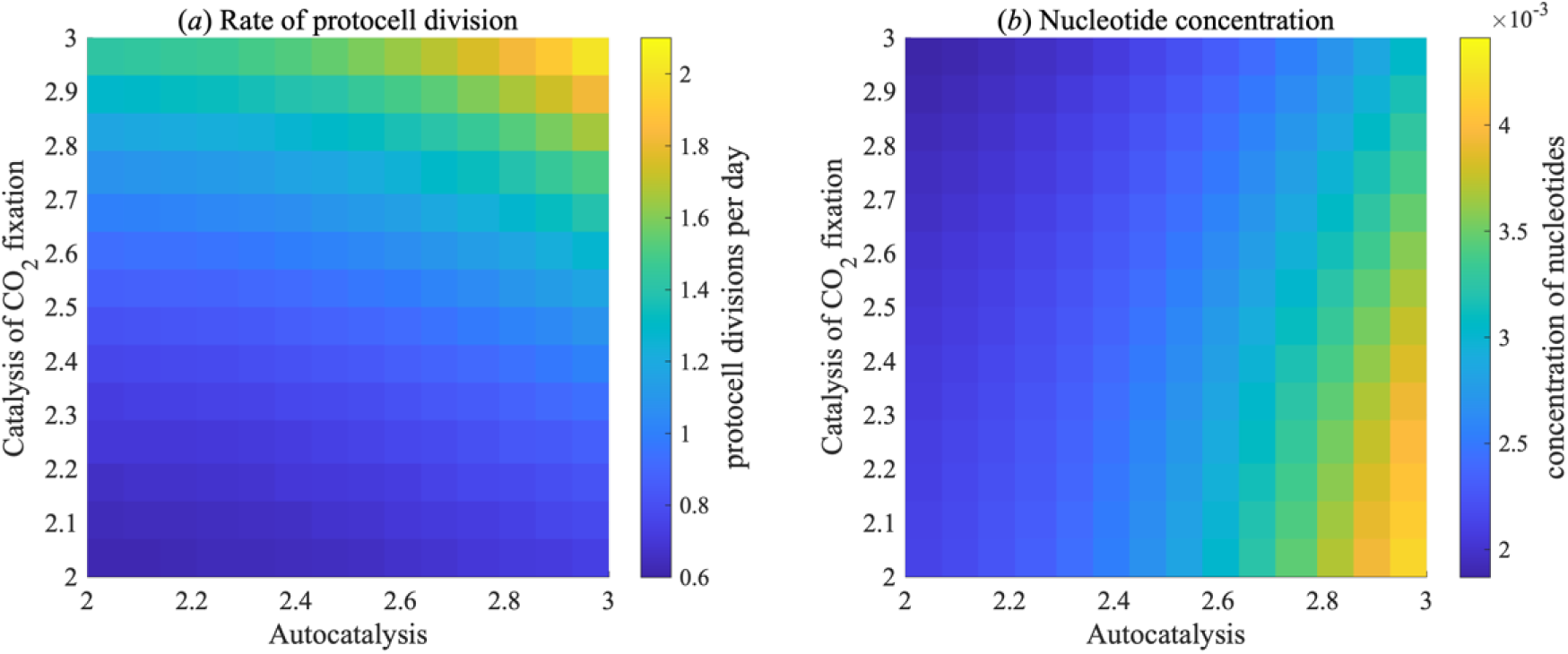
Impact of nucleotide autocatalysis and catalysis of CO_2_ fixation on protocell division rate and nucleotide concentration. Heat maps show (a) the number of protocell divisions per day and (b) the mean protocell concentration of nucleotides at equilibrium when varying the rate of nucleotide autocatalysis 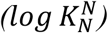 and nucleotide catalysis of CO_2_ fixation 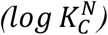. Other parameter values given in Table 1.

## Discussion

Prebiotic nucleotide synthesis is a necessary precursor for any form of RNA world. The model developed here explores how the catalytic activities of nucleotide monomers (‘naked cofactors’) could contribute to the autotrophic growth of protocells and the concentration of nucleotides inside them. Assuming the first protometabolic systems could at best generate trace amounts of nucleotides, nucleotide synthesis would need to be amplified through some form of positive feedback. We have analysed a range of different types of positive feedback, in which nucleotides catalyse CO_2_ fixation alongside a variety of branch points leading to the formation of particular molecular species (amino acids, fatty acids, sugars and a simple energy currency) as well as their own synthesis either in a direct (autocatalytic) or indirect manner (as a system outcome) (24, 57, 58). The results show that positive feedbacks can drive growth but only under certain topological limits on catalysis, some of which can also lead to the amplification of nucleotide concentration.

While the model deliberately simplifies protocell growth, our aim is to capture general features of metabolic heredity before the emergence of genes. The relative values of the reaction rate constants, specific concentrations and other parameters allow us to capture the topological network constraints on growth. The reactions and catalytic steps assumed in the model link directly with phylogenetic reconstructions of early cells as obligate chemiosmotic autotrophs that grew from the reduction of CO_2_ by H_2_ (42, 45, 73). The baseline proportion of carbon going into each metabolic branch is likewise based on thermodynamic estimates (11, 62). We find that the most important way in which nucleotides can promote protocell growth is by steepening the driving force that pushes protometabolic flux through the entire network. This is achieved by nucleotide catalysis of CO_2_ fixation, the conversion of CO_2_ and H_2_ into the reactive precursors of protometabolism (assumed to be prebiotic thioesters (59, 60); figure 1). The more these precursors form, the more they react to form the metabolic products needed for growth. Growth occurs directly when fatty acids are produced that contribute to the membrane, or indirectly through the production of amino acids (*AA*_1_) that chelate FeS clusters.

Reactive equivalents to the *C*_2_ precursor modelled here have been synthesised experimentally under relevant prebiotic conditions (50, 63). Likewise, fatty acids and amino acids are not only thermodynamically favoured (11, 62) but have been synthesised experimentally under relevant conditions (17, 22, 64-67). While sugars, notably ribose, have been synthesised under similar conditions (68, 69) they are less thermodynamically favoured (11, 62), so we assume that they are formed at arbitrarily low levels (5% of *C*_2_, Table 1). We also assume that acetyl phosphate (a plausible prebiotic energy currency (23, 47-49, 70, 71)) is formed at low levels, as the prebiotic availability of phosphate was probably limiting (52-56), and experiments achieved only low yields of acetyl phosphate under relevant conditions (48). Once formed, we assume that acetyl phosphate can phosphorylate molecules in a similar fashion to ATP (again demonstrated experimentally (48, 49)) for example in nucleotide synthesis, after which the acetate is lost as waste.

Another way in which nucleotides could promote protocell growth is by catalysing individual protometabolic pathways (figure 1*c*). This does not change the total flux through the system. It diverts flux down one pathway at the expense of others and adjusts the ratio of products formed away from the thermodynamically assigned proportions. When nucleotide catalysis of CO_2_ fixation is weak (figure 3*a*), additional catalysis of most pathways (fatty acids, energy currency or sugars) is neutral or detrimental. For example, making more fatty acids increases the protocell surface area but diverts flux away from amino-acid synthesis needed for CO_2_ fixation, which slows protocell growth. Catalysing the synthesis of an energy currency and sugars has little impact, as the baseline proportion of fixed carbon channelled into these products is low (Table 1), and their combination to form nucleotides yields little benefit; so increasing their production has little effect on protocell growth. The only useful target of downstream nucleotide catalysis is amino-acid synthesis, especially the amino acids that chelate FeS clusters (figure 3*a* and SI figure 1), as that directly promotes CO_2_ fixation. There is modest improvement in protocell growth with weak or moderate catalysis of amino-acid synthesis. But importantly, strong catalysis unbalances protometabolism, as it shifts too much carbon away from the fatty acids needed for the expansion of membrane surface area, collapsing protocell growth. This finding stresses the primacy of balanced flux for protocell growth, as catalysis that excessively diverts flux down one branch at the cost of others halts growth entirely.

When nucleotide catalysis of CO_2_ fixation is faster (figure 3*b*) the broad patterns are similar, except that now the synthesis of an energy currency becomes beneficial (figure 3*b*). Because a minimal proportion of fixed carbon is converted into energy currency, its availability limits nucleotide synthesis (Table 1). Making more energy currency therefore increases nucleotide synthesis, which in turn promotes CO_2_ fixation and protocell division. The cost of this positive feedback is negligible, as the channelling of carbon away from amino acids and fatty acids into an energy currency is trivial. The proportion of flux channelled into those pathways only falls slightly (because they were thermodynamically favoured in the first place). At the same time, overall protometabolic flux rises substantially through fast nucleotide catalysis of CO_2_ fixation. These findings assume no competition between nucleotide catalytic functions. Introducing a cost, in which the catalysis of specific branch points restricts nucleotide CO_2_ fixation (i.e. the same catalyst cannot do two things at once) undermines protocell division (figure 3*c*).

Nucleotide catalysis of specific pathways can also affect nucleotide concentration. Accelerated synthesis of nucleotide precursors (amino acids, sugars and energy currency) results in varying levels of nucleotide accumulation in protocells. Nucleotide concentration only increases sharply when the energy-currency pathway is favoured (figure 4), as energy is limiting; favouring this pathway feeds forward into a greater nucleotide synthesis. But nucleotide accumulation is lower when nucleotide catalysis of CO_2_ fixation is fast (figure 4*b*), as the increased number of nucleotides feeds back on protocell growth, speeding cell division and so halving nucleotide concentration more frequently. Greater nucleotide concentration can also occur through catalysis of amino acid synthesis (figure 4), but here for the illusory reason that strong catalysis collapses cell division (by undermining fatty acid production).

This important relationship between catalyst concentration and growth rate is emphasised by nucleotide autocatalysis. The prime example of this is purine synthesis, which relies on ATP, meaning ATP is needed to make ATP (72). If nucleotides catalyse CO_2_ fixation weakly, then autocatalysis contributes little to protocell growth, whatever the strength of autocatalysis (figure 5*a*, 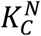 low, 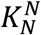 high). But with faster rates of CO_2_ fixation, cell division is strongly enhanced by autocatalysis (figure 5*a*, 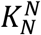 high, 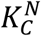 high). We anticipated that autocatalysis would generate excess nucleotides, but this is not the case in rapidly growing protocells (figure 5*b*). Instead, higher concentrations of nucleotides speed up protocell growth, lowering the nucleotide content as protocell division occurs more rapidly. This is an important general point: nucleotides will not accumulate in protocells if they have catalytic functions that speed up protocell growth and division. Thus, the premise for our model – protometabolic nucleotide synthesis requires catalytic positive feedbacks – precludes their rising concentration in actively growing protocells.

The model points to a second general constraint on nucleotide catalysis: only balanced protometabolism can sustain growth. Strong catalysis of specific pathways tends to unbalance protometabolism and collapse protocell growth. Conversely, non-specific catalysis of general processes that apply across the network – promiscuous catalysis – limits this problem. In fact, most universally conserved nucleotide cofactors are promiscuous, including NADH, FADH, acetyl CoA and ATP (42, 43, 45, 73). These promiscuous cofactors catalyse equivalent processes, such as the transfer of hydrogen, electrons, phosphate, acetyl or methyl groups (all fundamental features of metabolism), in many pathways across the entire metabolic network (3, 74, 75). Cofactors can catalyse these reactions without their enzymes (35-37), so the idea of an early era of biology when nucleotide monomers were the main catalysts has long held appeal (32, 33). Analysis of conserved biochemical pathways shows that NADH is indeed central to the earliest metabolic networks (57). Precisely because nucleotide cofactors are promiscuous, they do not excessively divert carbon flux down any particular branch. The pterins and folates derived from GTP are a rare example of non-promiscuous cofactors, as they specialise in C1 metabolism. But in fact these prove the rule, as they promote CO_2_ fixation, driving balanced flux through the whole network.

The properties of our model offer distinctive insights into the emergence of genetic selection. First, the model assumes that protometabolism occurs exclusively within growing protocells. Steepening the driving force or lowering kinetic barriers to overall flux will not promote growth if flux through any of the necessary protometabolic pathways is impeded by a low availability of externally sourced intermediates (‘food’). On the contrary, increased flux would unbalance metabolism and slow growth if some pathways are impeded. It is unlikely that nucleotide cofactors as complex as NADH and ATP could be reliably sourced from the environment, so this assumption seems reasonable. Thus, protocell metabolism can regenerate itself only if it is a system property of the protocell, where catalytic nucleotides are physically inherited at cell division through direct metabolic heredity. If indeed the core of metabolism can arise and reproduce itself spontaneously in this way, improving over time through positive feedbacks, then the later evolution of genetically encoded catalysts (enzymes and ribozymes) becomes less of a formidable problem. Sequences that enhance protometabolic flux, either directly as ribozymes or indirectly as polypeptides that are templated through biophysical interactions between amino acids and RNA sequences, will promote protocell growth through exactly the same types of catalytic feedback explored here. In contrast, those that act selfishly or interfere with protocell growth will be selected against.

Second the model shows that weak catalysis of specific pathways is better than strong catalysis, while promiscuous broad-spectrum processes are better than more focused ‘enzymatic’ catalysts. Weak catalysis does not unbalance metabolism because it only marginally diverts flux down specific pathways at the expense of others, whereas strong catalysis can collapse the system by diverting flux away from essential pathways (figure 3). Promiscuous processes such as hydrogen transfer do not unbalance metabolism, but are intrinsically less sophisticated than enzyme catalysis; similar processes occur with FeS clusters, metal ions or minerals, giving a seamless transition from geochemistry to biochemistry (41). This finding mirrors previous modelling work showing that, unlike modern phospholipid membranes, rudimentary bilayers made from fatty acids can freely exchange ions, allowing hydrothermal flow to maintain disequilibria (such as pH gradients) without the need for sophisticated pumps (77, 78). For autotrophic growth, only gases such as H_2_, CO_2_, NH_3_ and HS^-^ are needed, along with metal ions and protons, all of which readily cross leaky fatty-acid membranes (78). The idea that simplicity precedes complexity is not just pleasing but necessary, for only simplicity works.

In conclusion, the model developed here shows there are constraints on how flux must operate in protocells. Catalysts must facilitate balanced flux through protometabolism to promote fast protocell growth via positive feedbacks. Selection for fast growth is likely to oppose the accumulation of catalytic nucleotides within protocells, albeit positive feedbacks favouring the synthesis of an energy currency increase nucleotide concentrations somewhat. Most importantly, the model gives a new context for the emergence of the genetic code in autotrophic protocells. Whatever processes allowed polymerization of nucleotides in this setting, the introduction of short, random RNA sequences inside replicating protocells offers an immediate informational context. Rather than ‘inventing’ information from nothing, the first peptides or ribozymes would drive protocell growth through interactions with cofactors, promoting balanced flux through the same branching network that is still conserved today. So protocell growth gives meaning to information from the origins of polymerization, laying the foundations for the emergence of the genetic code.

## Acknowledgements

This work was supported by funding from the Engineering and Physical Sciences Research Council (EP/F500351/1, EP/I017909/1) and Natural Environment Research Council (NE/R010579/1) to AP, the Biotechnology and Biological Sciences Research Council (BB/S003681/1) and bgc3 to NL, and a joint grant to AP and NL from the Biotechnology and Biological Sciences Research Council (BB/V003542/1).

