## Supplementary Information for "The limits of metabolic heredity in protocells"

**Table S1:** Other parameters used for simulations.

| Parameter | Symbol | Value | Unit |
|---|---|---|---|
| Saturating constant for the catalysis of carbon fixation by amino acids (type 1) | $Km_{AA_1}$ | $10^{-4}$ | $mol\ dm^{-3}$ |
| Rate constant for carbon fixation | $K^{C_2}$ | $10^5$ | $mol^{-1}\ dm^3 s^{-s}$ |
| Saturating constant for catalysis of nucleotide production by nucleotides | $Km_N$ | 2 | $mol\ dm^{-3}$ |
| Permeability constant for all species | $\phi$ | $10^{10}$ | $cm\ s^{-1}$ |
| Stoichiometry constant for synthesis of fatty acids from $C_2$ | $v_{FA}$ | 5 | – |
| Stoichiometry constant for synthesis of amino acids from $C_2$ | $v_{AA}$ | 1 | – |
| Stoichiometry constant for synthesis of energy from $C_2$ | $v_E$ | 1 | – |
| Stoichiometry constant for synthesis of sugars from $C_2$ | $v_S$ | 3 | – |

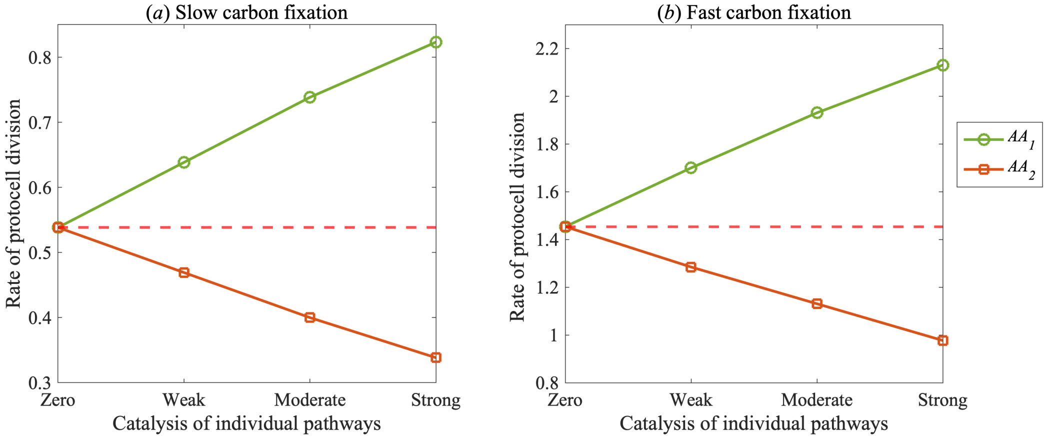


**Figure S1: Impact of nucleotide catalysis of amino acid types on protocell division rate.** Changes in cell division per day with nucleotide catalysis of the synthesis pathways of amino acids that chelate crystals to facilitate carbon fixation($AA_1$, green), and amino acids used to make nucleotides ($AA_2$, orange), given no ($K_i^N = 0$), weak ($K_i^N = 0.33$), medium ($K_i^N = 0.66$) and strong catalysis ($K_i^N = 0.99$) ($i = AA, FA, S, E$). This is shown under the assumption that $CO_2$ fixation is (a) slow ($K_{C_2}^N = 10^{0.9}$)or (b) fast ($K_{C_2}^N = 10^4$). The red dotted line indicates the number of protocell divisions per day with nucleotide catalysis of $CO_2$ fixation only (no catalysis of other synthesis pathways). Other parameter values are given in Table 1.

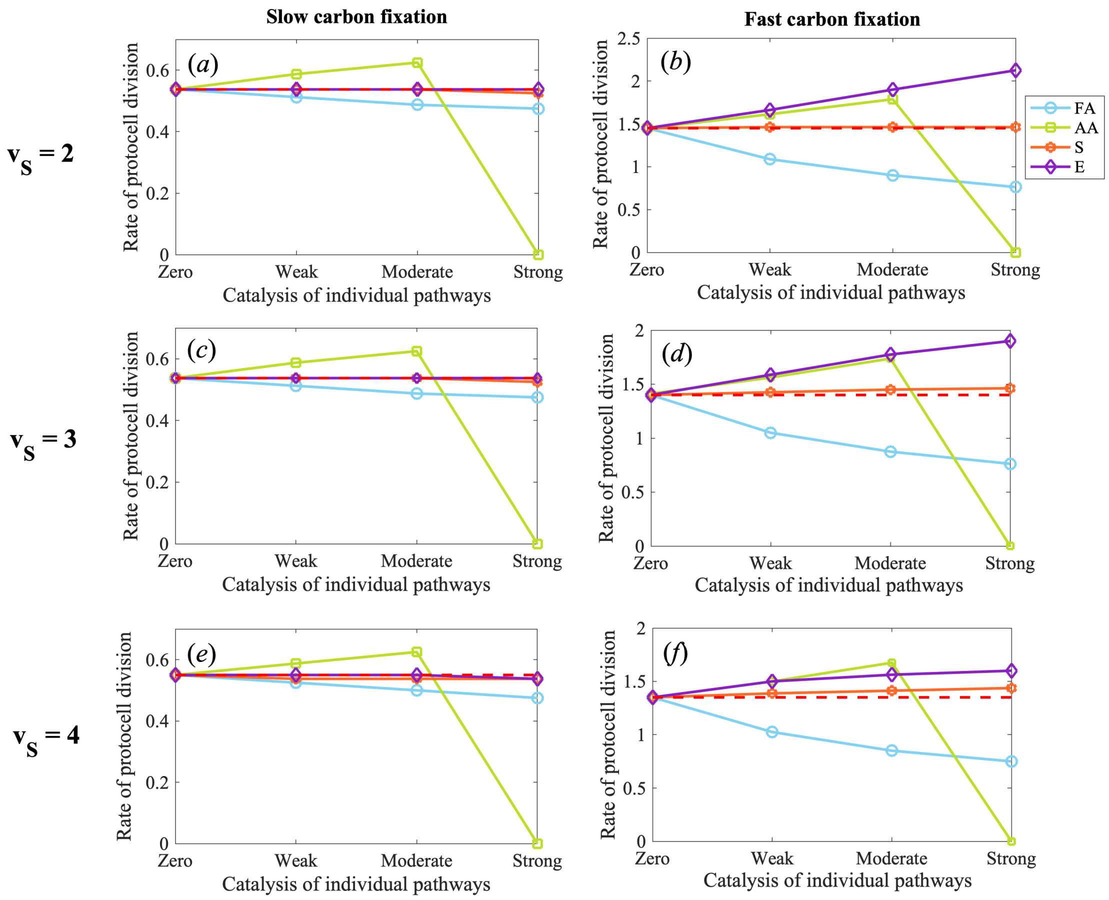


**Figure S2: Impact of nucleotide catalysis on protocell rate with different stoichiometric coefficients for the synthesis of sugars.** Changes in cell division per day with increasing rates of nucleotide catalysis of the synthesis pathways of amino acids (AA, green), fatty acids (FA, blue), energy (E, purple) and sugars (S, yellow), for increasing numbers of $C_2$ needed to make one molecule of sugar ($v_S$). This is shown on the x-axis for no catalysis ($K_i^N = 0$), weak catalysis ($K_i^N = 0.33$), medium catalysis ($K_i^N = 0.66$) and strong catalysis ($K_i^N = 0.99$), where $i = AA, FA, S, E$. This is shown under the assumption that $CO_2$ fixation is (a) slow ($K_{C_2}^N = 10^{0.9}$)or (b) fast ($K_{C_2}^N = 10^4$). The red dotted line indicates the number of protocell divisions per day with nucleotide catalysis of $CO_2$ fixation only (no catalysis of other synthesis pathways). Other parameter values are given in Table S1.

### S3: Model Structure

Species symbols and initial concentrations are shown in Table 1. The number of molecules each species present is denoted by the letter n followed by the species symbol in superscript (e.g. $n^S$ is the number of sugar molecules). Concentrations of different species are denoted by the species symbol followed by a location (cell or sink) in superscript (i.e. the concentration of Sugars in the cell is denoted by $S^{cell}$).

Table S2: Initial concentrations of chemical species in the protocell (all in $mol\ dm^{-3}$).

| **Chemical species** | **Symbol** | **Value** |
|---|---|---|
| Concentration of the two carbon reverse citric acid cycle intermediate in the cell | $C_2^{cell}$ | 0 |
| Concentration of the two carbon primitive energy currency (such as acetyl phosphate) in the cell | $E^{cell}$ | 0 |
| Concentration of fatty acids in the cell | $FA^{cell}$ | 0.0198 |
| Concentration of amino acids (both types) in the cell | $AA^{cell}$ | 0.0198 |
| Concentration of sugars in the cell | $S^{cell}$ | 0 |
| Concentration of nucleotides in the cell | $N^{cell}$ | 0 |
| Concentration of $CO_2$ in the sink | $CO_2^{sink}$ | 0.001 |
| Concentration of amino acids in the sink | $AA^{sink}$ | $10^{-6}$ |
| Concentration of primitive energy currency (E) in the sink | $E^{sink}$ | $10^{-6}$ |
| Concentration of nucleotides in the sink | $N^{sink}$ | 0 |
| Concentration of sugars in the sink | $S^{sink}$ | 0 |

**Cell volume and surface area** The protocell is assumed to be spherical and its volume ($V$) and surface area ($SA$) are functions of the number of fatty acids in the system, as fatty acids are assumed to instantly embed in the membrane. The protocell surface area is given by the number of fatty acids multiplied by the area of the fatty acid head group a:

$$SA = n^{FA}a \tag{1}$$

And the protocell volume is a spherical volume calculated with the radius given by the cell surface area:

$$V = (n^{FA}a)/3\sqrt{(n^{FA}a)/4\pi} \tag{2}$$

**Concentrations** The concentration of a species in the protocell is calculated using the following equation, where $A_N$ is Avogadro's number and $X$ is an arbitrary chemical species:

$$X^{cell} = n^X/A_N V \tag{3}$$

**Carbon fixation** In this model, carbon fixation is the synthesis of a two-carbon reverse citric acid cycle intermediate $C_2$. When only catalysed by amino acids, the synthesis of $C_2$ is directly proportional to the concentration of $AA_1$, with a rate constant $K_{C_2}$, and to the concentration of $CO_2$ in the sink via a saturating Michaelis-Menten equation with saturating constant ($Km_{AA_1}$):

$$\frac{dn^{C_2}}{dt} = AA_1 \frac{CO_2^{sink}}{CO_2^{sink} + Km_{AA_1}} K_{C_2} \tag{4}$$

In the formulation of the model where nucleotide co-factors can catalyse carbon fixation, the equation for synthesis of $C_2$ is as follows:

$$\frac{dn^{C_2}}{dt} = AA_1 \frac{CO_2^{sink}}{CO_2^{sink} + Km_{AA_1}} K_{C_2}(1 + K_{C_2}^N N^{cell}) \tag{5}$$

We assume that catalysis of individual pathways and catalysis of carbon fixation are done by different nucleotide species of equal concentrations - so there would be no cost one type of catalysis when introducing another. An alternative assumption was considered in Figure 3: that the same nucleotides perform both functions, so increasing the rate of one type of catalysis decreases the rate of the other. This was done by multiplying the one of the rate equations by a proportion $\alpha$ and the other by $1 - \alpha$.

$$\frac{dn^{C_2}}{dt} = AA_1 \frac{CO_2^{sink}}{CO_2^{sink} + Km_{AA_1}} K_{C_2}(1 + (1 - K_i^N) K_{C_2}^N N^{cell}) \tag{6}$$

**Synthesis of organics from $C_2$** Since $C_2$ is assumed to be instantly converted into another organic species. The expression for the rate of synthesis of the 5 organic molecules made directly from $C_2$ (fatty acids, both types of amino acids, sugars and $E$) is simply a proportion ($\lambda^{C_2}$) of the number of $C_2$ molecules in the protocell divided by a stoichiometry constant ($v_X$) (which depends on the number of C atoms used to make that molecule) :

$$\frac{dn^X}{dt}(from\ C_2) = \frac{\lambda_X^{C_2}\ n^{C_2}}{v_X} \tag{7}$$

**Nucleotide synthesis** Nucleotides are assumed to be made from equal parts of sugars and type 2 amino acids, and to have its synthesis facilitated by the primitive energy currency $E$. Therefore, the rate of synthesis of nucleotides is given by the probability of sugars, amino acids type 2 and $E$ molecules reacting (which is proportional to the concentration of the three species with a proportionality constant ($K_N^E$)) multiplied by the number of molecules of the limiting reagent .

$$\frac{dn^N}{dt}(byE) = K_N^E \; S^{cell} \; AA_1^{cell} \; E^{cell} \; n^{lim} \tag{8}$$

Equation 9 describes nucleotide synthesis when nucleotides catalyse their own production. A Michaelis-Menten expression is used to account for the fact that, unlike $E$, nucleotides do not get used up as they react. Instead there is a period of time (i.e. between being reduced and oxidised again) that nucleotide molecules cannot be used in reactions.

$$\frac{dnN}{dt}(byN) = K_N^N \; \frac{S^{cell}}{S^{cell} + Km_N} \; \frac{AA_1^{cell}}{AA_1^{cell} + Km_N} \; N^{cell} \; n^{lim} \tag{9}$$

**Nucleotide catalysis of individual synthesis pathways** Where the possibility of nucleotides catalysing the formation of different organic molecules (from $C_2$) was explored, this process was modelled by increasing the proportion of $C_2$ that is used to produce each species ($\lambda C_2$) proportionally to the concentration of nucleotides in the protocell, using a Michaelis-Menten expression with a saturating constant $Km_N$. As the proportion of $C_2$ being used up in the production of one molecule increases, the proportion of $C_2$ used in producing all other molecules decreases accordingly. The proportion between rates of production of the uncatalysed species is maintained, so the ratios remain accordant with thermodynamic data [32].

$$\lambda_i^{C_2} = \lambda_i^{C_2} \; (1 + K_i^N \; \frac{N^{cell}}{N^{cell} + Km_N}) \tag{10}$$

We assume that catalysis of individual pathways and catalysis of carbon fixation are done by different nucleotide species of equal concentrations - so there would be no cost one type of catalysis when introducing another. An alternative assumption was considered in Figure 3: that the same nucleotides perform both functions, so increasing the rate of one type of catalysis decreases the rate of the other. This was done by multiplying the one of the rate equations by a proportion $\alpha$ and the other by $1 - \alpha$.

$$\frac{dn^{C_2}}{dt} = AA_1 \frac{CO_2^{sink}}{CO_2^{sink} + Km_{AA_1}} K_{C_2}(1 + (1 - \alpha)K_{C_2}^N N^{cell}) \tag{11}$$

**Loss by diffusion** The total change in the number of molecules of a species inside the protocell due to diffusion is given by the concentration gradient between sink and cell multiplied by the surface-area to volume ratio and a permeability constant ($\phi$)(Equation 12). Both types of amino acids, sugars, E and nucleotides are subject to gain or loss by diffusion; $C_2$ and fatty acids are not subject to diffusion because the former gets instantly converted into other organic molecules, while the latter gets instantly embedded in the membrane.

$$\frac{dn^X}{dt}(diffusion) = (X^{sink} - X^{cell}) \; \frac{SA}{V} \; \phi \; dt \tag{12}$$

**Fatty acid loss at division** Loss of fatty acids molecules at division was simply done by subtracting a percentage of of fatty acids from the protocell before division - so the "daughter cell" inherits half of the resultant number of fatty acids.

**Total change in number of molecule of each species in the protocell** The total change in the number of molecules of each species is given by their synthesis, gain or loss by diffusion and loss by being a reactant in the formation of another molecule.

$$\frac{dn^{C_2}}{dt}(total) = \frac{dn^{C_2}}{dt} - \frac{dn^{E}}{dt} - \frac{dn^{FA}}{dt} - \frac{dn^{AA_1}}{dt} - \frac{dn^{AA_2}}{dt} \tag{13a}$$

$$\frac{dn^{E}}{dt}(total) = \frac{dn^{E}}{dt} - \frac{dn^{N}}{dt}(byE) + \frac{dn^{E}}{dt}(diffusion) \tag{13b}$$

$$\frac{dn^{FA}}{dt}(total) = \frac{dn^{FA}}{dt} \tag{13c}$$

$$\frac{dn^{AA_1}}{dt}(total) = \frac{dn^{AA_1}}{dt} + \frac{dn^{AA_1}}{dt}(diffusion) \tag{13d}$$

$$\frac{dn^{AA_2}}{dt}(total) = \frac{dn^{AA_2}}{dt} - \frac{dn^{N}}{dt} + \frac{dn^{AA_2}}{dt}(diffusion) \tag{13e}$$

$$\frac{dn^{S}}{dt}(total) = \frac{dn^{S}}{dt} - \frac{dn^{N}}{dt} + \frac{dn^{S}}{dt}(diffusion) \tag{13f}$$

$$\frac{dn^{N}}{dt}(total) = \frac{dn^{N}}{dt}(byE) - \frac{dn^{N}}{dt}(byN) + \frac{dn^{N}}{dt}(diffusion) \tag{13g}$$

**Integration of dynamic equations** A single-step forward Euler integration was used with a step size of 1 sec. At each time step the number of molecules in the protocell was rounded to the nearest integer, since there cannot be a fraction of a molecule.

**Cell division** Protocells divide when they reach a critical surface area size (measured by proxy as the number of fatty acid molecules), and each daughter cell inherits half of the contents of the mother cell. In order to evaluate the robustness of the model, we also considered a stochastic mechanism of inheritance (see next paragraph).

**Parameters and Matlab script** This model was implemented using Matlab (vR2018b, The Mathworks, MA). The Matlab scripts are available in a repository on GitHub (`https://github.com/raquelnpalmeira/limits_of_metabolic_heredity`). Initial conditions and parameters used for simulations can be found in Tables 1 and S1 (where parameters vary from the values in the tables, they are set out explicitly in the text).
